# Peculiar cases of a “sleeping” brain in alert cancer patients

**DOI:** 10.1101/732230

**Authors:** Helene Benveniste, Paul Vaska, Dinko Franceschi, Michael Salerno, Sabeen Rizwan, Hedok Lee, Jean Logan, Douglas Rothman, Yuri Lazebnik, Nora D. Volkow, Thomas V. Bilfinger

## Abstract

Cognitive and constitutional symptomatology is common in cancer patients but the causes are not well understood. To investigate whether cancers cause these symptoms by changing cerebral metabolism, we measured the cerebral rate of glucose consumption (CMRglc) in patients diagnosed with a lung lesion.

**Methods:** The CMRglc was quantified in 20 patients undergoing ^18^F-FDG PET for lesion staging. The cognitive status was assessed by neuropsychological testing.

**Results:** Paradoxically, despite being alert three of the patients had CMRglc typical of people who are in deep sleep or anesthetized. All three had cancers, two died within 2 months of scanning. Remaining patients including four with early stage cancer had CMRglc within normal range.

**Conclusions:** We speculate that the low CMRglc reflects a switch to an alternative energy source that is mediated by cancers remotely. Identifying the underlying mechanism and the alternative energy sources may help to understand how cancers can change body metabolism.

## Introduction

Lung cancer is often diagnosed at an advanced stage when the majority of patients already have constitutional and central nervous system (CNS) symptomatology, including fatigue and sleep perturbations, but how cancers cause these debilitating symptoms is poorly understood. One possibility is that cancers can affect CNS remotely. It is known that a cancer residing in one organ can secrete signaling molecules, such as cytokines that remotely affect the functions of other organs, including the brain. In a retrospective study, patients with lung cancers had altered glucose uptake (standard uptake values, SUV) in various regions of the brain (*1*) but whether these changes could cause CNS dysfunction is unknown. Moreover, assessing brain function requires measuring the absolute cerebral metabolic rate of glucose consumption (CMRglc), which SUV does not provide.

Our hypothesis was that the CNS dysfunction would correlate with low brain function and thus lower CMRglc. To quantify brain function, we measured the CMRglc in patients diagnosed with lung nodules while they were undergoing whole-body ^18^F-FDG-PET for staging of the lung lesions. CMRglc was calculated by combining this procedure with measurement of ^18^F time-activity in blood (*2*).

## Materials and methods

The research protocol was approved by the Institutional Review Board at Stony Brook University and written informed consent was obtained from each patient. Inclusion criteria: 1) >18 years of age; 2) fluency in English; 3) patient with a lung nodule referred for FDG PET-CT scan. Exclusion criteria: 1) chemo-and/or radiation therapy within 6 months prior; 2) hearing or vision impairment that interfere with neuropsychological testing; 3) significant dementia (not able to sign consent for clinical treatment);4) history of severe psychiatric illness; 5) history of or current dialysis treatment; 6) under treatment for active hepatitis; 7) clinically unmanaged diabetes; 8) treatment with cognitively stimulating medications. The data from twenty healthy subjects reported previously (*3*) were used as controls.

### Neuropsychological testing

The patients also underwent a psychometric test battery to assess their fatigue, cognitive function, mood status and pain immediately prior to PET scanning (Supplemental Box 1 and Table 1S).

### ^18^FDG PET procedures

The patients fasted overnight. One venous catheter was inserted in a dorsal hand vein which was kept warm to arterialize the venous blood for sampling (*4*). Patients received 10–15 mCi of ^18^F-FDG intravenously and instructed to relax to obtain CMRglc under resting conditions. The time-course of the arterialized-venous plasma ^18^F activity was determined and corrected for radioactive decay to the scan start time (*2*). Plasma glucose concentrations was measured 4 times during the study.

PET scans were conducted with a Biograph Truepoint 40 PET-CT scanner (Siemens Medical Solutions USA Inc., Malvern, Pennsylvania, USA) with a 40-detector-row helical CT scanner. Scanning started 40-min after ^18^F-FDG injection and lasted for 20 minutes. Attenuation maps were constructed using the CT data. The PET data were reconstructed with the vendor’s ordered-subset expectation maximization algorithm. The resulting images had a voxel size of 1.02×1.02×1.5 mm and were smoothed with an isotropic 2-mm FWHM Gaussian filter.

The FDG scans were transformed into metabolic images using the MRGlu (FDG Autorad) quantification methods implemented in PMOD (PMOD 3.6 LLC, Zurich Switzerland) based on an autoradiographic solution model (*5*). The following parameters were used for the CMRglc calculation: Lumped constant=0.52; K1=0.095; k2=0.125; k3=0.069 and k4=0.0055 (*2*). The maximum probability atlas (Hammers N30R83) was used for VOI definition. Global CMRglc values were calculated by a volume-weighted average of all VOI’s defined including the cerebellum and brainstem.

## Results

We have found that CMRglc in three of the patients was about 20% lower than in others (**Table 1 and Fig. 1**) and in the range reported for anesthesia (*6*), deep wave sleep (*7*), Alzheimer’s dementia (*8*), locked-in state (*9*), or stroke (*10*). In brain areas that normally consume more glucose than others (occipital lobe), the CMRglc was reduced by ∼30% (**Fig. 2**). Moreover, when compared to 20 normal subjects who were analyzed previously (*3*), the CMRglc decrease of the three patients (**Fig. 1**) approached the 42% reduction reported for coma patients (*11*). Thus, according to common knowledge, the brain function of the three patients had to be near the threshold beyond which conscious activity cannot be maintained (*11*). Yet, the three patients were alert on the day of the PET procedure and passed psychometric tests immediately before the scanning with no signs of major cognitive impairment, excessive pain, or fatigue (Supplemental Table 1S).

**Figure 1:**
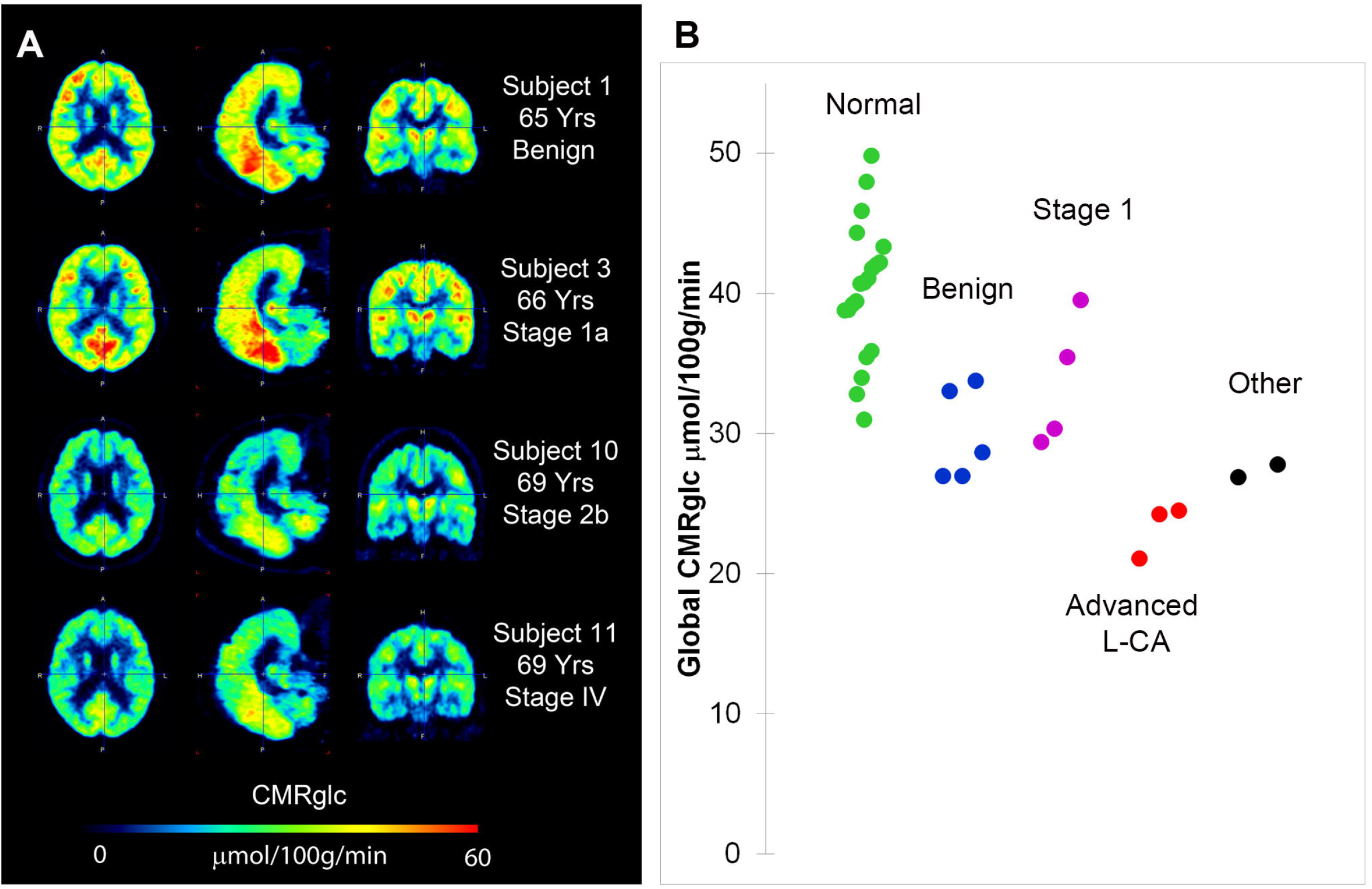
**A:** Cerebral metabolism rate of glucose consumption (CMRglc) scans from two subjects with advanced stage lung cancer with abnormally low CMRglc compared to a subject with a benign lung lesion and a subject with early stage lung cancer. **B:** Global CMRglc of the 20 study patients in comparison to 20 healthy individuals. Each dot represents the global brain CMRglc for one individual. Green: Normal subjects (N=20, Age: 19-51 yrs.); Blue: Patients with benign lung nodules (N=5, 62-75 yrs.); Magenta: Patients with non-small cell lung cancer (NSCLC) Stage 1 (N=5, 48-65 yrs.); Red: Patients with advanced NSCLC lung cancer (N=3, 51-69 yrs.); Black: Patients with other cancers (N=2, 72 and 65 yrs.).

**Figure 2:**
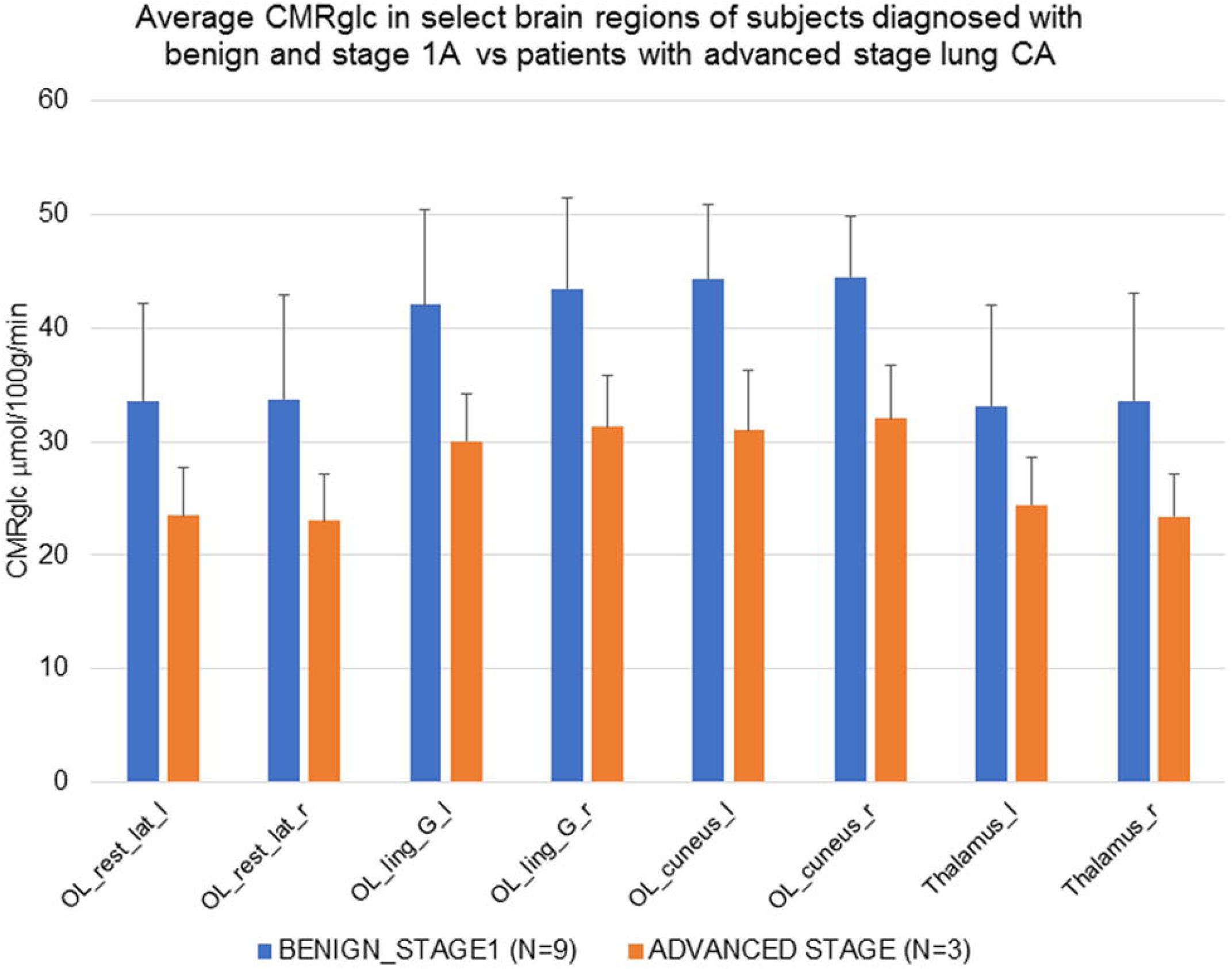
The CMRglc in brain regions of subjects with benign lesions and stage 1 lesions (blue) in comparison to the three patients with advanced stage lung cancer. The CMRglc of the occipital lobe (OL) is approximately 20 % lower in the patients with advanced stage lung cancer. OL_rest_lat_l=Occipital lobe lateral remainder of occipital lobe (Left); OL_rest_lat_r=Occipital lobe lateral remainder of occipital lobe (Right); OL_ling_G_l=Occipital Lobe Lingual Gyrus (Left); OL_ling_G_r=Occipital Lobe Lingual Gyrus (Right); OL_cuneus_l=Occipital Lobe Cuneus (Left); OL_cuneus_r=Occipital Lobe Cuneus (Right); Thalamus_l=Thalamus (Left); Thalamus_r=Thalamus (Right).Disclosure: The authors declare no conflicts of interest.

**Table 1:**
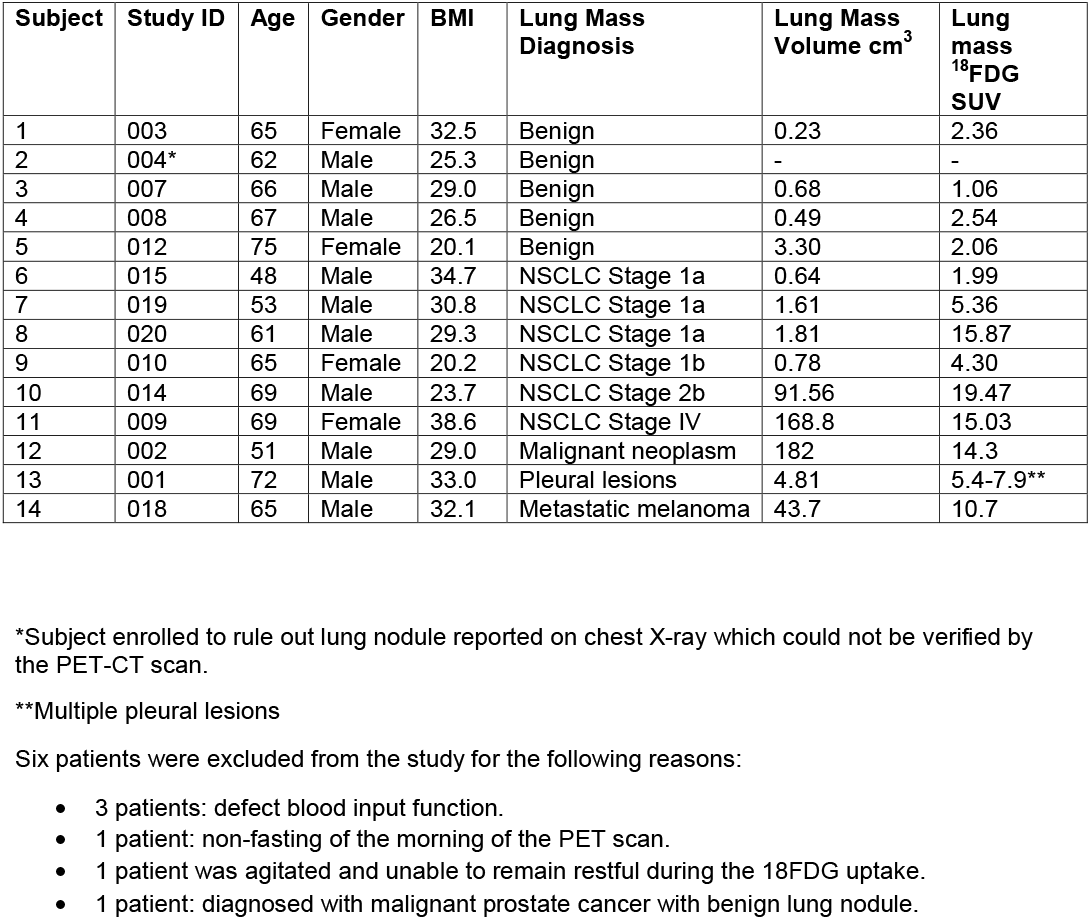
Age, gender and lung nodule diagnosis

| Subject | Study ID | Age | Gender | BMI | Lung Mass Diagnosis | Lung Mass Volume cm<sup>3</sup> | Lung mass <sup>18</sup>FDG SUV |
| --- | --- | --- | --- | --- | --- | --- | --- |
| 1 | 003 | 65 | Female | 32.5 | Benign | 0.23 | 2.36 |
| 2 | 004* | 62 | Male | 25.3 | Benign | - | - |
| 3 | 007 | 66 | Male | 29.0 | Benign | 0.68 | 1.06 |
| 4 | 008 | 67 | Male | 26.5 | Benign | 0.49 | 2.54 |
| 5 | 012 | 75 | Female | 20.1 | Benign | 3.30 | 2.06 |
| 6 | 015 | 48 | Male | 34.7 | NSCLC Stage 1a | 0.64 | 1.99 |
| 7 | 019 | 53 | Male | 30.8 | NSCLC Stage 1a | 1.61 | 5.36 |
| 8 | 020 | 61 | Male | 29.3 | NSCLC Stage 1a | 1.81 | 15.87 |
| 9 | 010 | 65 | Female | 20.2 | NSCLC Stage 1b | 0.78 | 4.30 |
| 10 | 014 | 69 | Male | 23.7 | NSCLC Stage 2b | 91.56 | 19.47 |
| 11 | 009 | 69 | Female | 38.6 | NSCLC Stage IV | 168.8 | 15.03 |
| 12 | 002 | 51 | Male | 29.0 | Malignant neoplasm | 182 | 14.3 |
| 13 | 001 | 72 | Male | 33.0 | Pleural lesions | 4.81 | 5.4-7.9** |
| 14 | 018 | 65 | Male | 32.1 | Metastatic melanoma | 43.7 | 10.7 |

\*Subject enrolled to rule out lung nodule reported on chest X-ray which could not be verified by the PET-CT scan.
\*\*Multiple pleural lesions
Six patients were excluded from the study for the following reasons:
- 3 patients: defect blood input function. - 1 patient: non-fasting of the morning of the PET scan. - 1 patient was agitated and unable to remain restful during the <sup>18</sup>FDG uptake. - 1 patient: diagnosed with malignant prostate cancer with benign lung nodule.

To determine whether this paradoxical observation was a technical artifact, we verified all parameters and procedures used to calculate CMRglc in each subject (Supplemental Fig. 1S). We found no errors that could account for our observation. We therefore concluded that the calculated CMRglc in all patients faithfully represented the actual glucose consumption.

## Discussion

In this study, we documented that in some cancer patients the cerebral rate of glucose consumption (CMRglc) can be as low as in individuals who are asleep, under anesthesia or comatose. Nevertheless, these patients were alert and functioned normally, as indicated by the results of neurocognitive tests. We propose that the simplest explanation of this phenomenon is that their brains had switched from glucose to an alternative fuel.

That the brain can switch to an alternative fuel under some circumstances is well known. Glucose can be replaced by lactate during extensive physical exercise (*12*), by ketones during starvation (*13*) or by acetate during alcohol intoxication (*14*). The switch to ketones or some other mono carboxylic acids (*15*) can support up to 50% of the brain energy demands, which in principle, could explain how the cancer patients functioned cognitively despite the low CMRglc. However, none of the conditions known to us that can switch energy consumption to lactate or ketones were applicable in our case, as the patients were resting, not intoxicated, were not heavy drinkers by history, and maintained a regular diet (Supplemental Table 2S).

Because advanced cancer was the only common condition that we identified in the three patients, we speculate that the metabolic switch was a result of malignancy and suggest calling it ‘**C**ancer **A**ssociated **E**nergy **S**witch’, or CAES. The CAES hypothesis raises several questions.

How does cancer cause CAES? Because the brain scans of the CAES patients showed no signs of neoplasia, the effect of cancer on CMRglc had to be remote. Given the heterogeneity of malignant lesions and the ability of cancer cells to ectopically secrete a variety of regulatory molecules, including cytokines, it is plausible that some of them can induce CAES. Identifying these molecules will require a systematic investigation, which we hope our report will encourage. The candidate molecules should explain how they could change glucose metabolism in the brain without changing it systemically, as the blood glucose of the patients with CAES was normal.

What is the energy substrate(s) that is used in CAES instead of glucose? In principle, it can be a substitute known to be used by the brain, such as ketones, lactate, or glutamine, or other, yet to be identified, candidates.

How common is CAES in cancer? We found this phenomenon by analyzing a group of lung cancer patients, but whether it is present in other cancers remains to be determined.

Given that FDG-PET is a routine procedure in cancer screening and in measuring activity of brain regions, an obvious question is: Why were patients with CAES overlooked previously? We suggest that the answer is in how glucose consumption is measured. The routine diagnostic FDG-PET measures *relative* rates of glucose consumption (SUV), which is sufficient to detect cancers because they consume more glucose than the surrounding normal tissue. Likewise, measuring SUV of glucose uptake is sufficient to compare the activities of different brain area. However, to compare the rate of glucose consumption across patients requires measuring CMRglc, because it represents glucose consumption in *absolute* units (μmol/100g/min). This is what we did in our study which allowed us to make the paradoxical observation that we report.

In summary, we provide evidence consistent with the hypothesis that malignancy can affect brain energy metabolism remotely, the phenomenon we named ‘**C**ancer **A**ssociated **E**nergy **S**witch’, or CAES. We suggest that understanding the underlying mechanisms may help to alleviate systemic effects of cancer.

