## Supplemental Table 1 for "Peculiar cases of a “sleeping” brain in alert cancer patients"

**Table 1S: Neurocognitive, pain, fatigue and mood status of the enrolled patients**

| Subject | Lung mass diagnosis | Study ID | FACIT-F (% of Total) | SDMT (SD)* | CESD | NRS | Karnofsky | WTARVIQ |
|---|---|---|---|---|---|---|---|---|
| 1 | Benign | 003 | 0.93 | 41 (-0.25) | 8 | 0 | 100 | 121 |
| 2 | Benign | 004 | 0.61 | 43 (-0.5) | 12 | 7 | 60 | 107 |
| 3 | Benign | 007 | 0.97 | 35 (-0.75) | 0 | 8 | 90 | 108 |
| 4 | Benign | 008 | 0.89 | 38 (-0.5) | 6 | - | 100 | 114 |
| 5 | Benign | 012 | 0.87 | 35 (-0.75) | 15 | 0 | 100 | 100 |
| 6 | NSCLC Stage 1a | 015 | 0.55 | 39 (-0.75) | 35 | 3 | 90 | 99 |
| 7 | NSCLC Stage 1a | 019 | 0.87 | 46 (-0.25) | 1 | 2 | 90 | 119 |
| 8 | NSCLC Stage 1a | 020 | 0.94 | 44 (0) | 1 | 0 | 100 | 99 |
| 9 | NSCLC Stage 1b | 010 | 0.56 | 40 (-0.25) | 46 | 2 | 90 | 107 |
| 10 | NSCLC Stage 2b | 014 | 0.70 | 29 (-1.5) | 4 | 5 | 80 | 103 |
| 11 | NSCLC Stage IV | 009 | 0.67 | 43 (1) | 21 | 8 | 80 | 107 |
| 12 | Malignant neoplasm | 002 | 0.77 | 33 (-1.5) | 18 | 6 | 80 | 111 |
| 13 | Pleural malignancy | 001 | 0.99 | 38 (0.5) | 3 | 3 | 90 | - |
| 14 | Metastatic melanoma | 018 | 0.83 | 48 (0.5) | 14 | 0 | 90 | 118 |

*Presented as raw scores.

The three patients with decreased CMRglc are highlighted.
