## Supplemental Table 2 for "Peculiar cases of a “sleeping” brain in alert cancer patients"

**Table 2S: Comorbidity and medications**

| Subject # (Study ID) | Hypercholesterolemia or Hyperlipidemia | Hypertension | CAD | Depression | Chronic Pain | Asthma | Diabetes | Alcohol Use | Tobacco Use (pack yrs) |
|---|---|---|---|---|---|---|---|---|---|
| 1 (003) | - | Atenolol | - | - | - | - | - | 2/month | 12.5 |
| 2 (004) | Crestor | Metoprolol, Lisinopril | - | - | Dilaudid, Tramadol, baclofen, ASA 325 | - | - | 2/week | 0 |
| 3 (007) | Crestor | - | - | - | Tegretol | - | - | 7/week | 75 |
| 4 (008) | - | - | - | - | Flexeril, Tramadol, Vicodin | - | - | 0 | 82.5 |
| 5 (012) | - | - | - | - | - | - | | 6/month | 15.375 |
| 6 (015) | Crestor | Carvedilol, Lisinopril | Plavix | Zoloft | - | - | - | 2/month | 40 |
| 7 (019) | - | - | - | - | - | symbicort, albuterol, Singulair | - | 0 | 25 |
| 8 (020) | Zocor | Norvasc, HCTZ | Toprol XL, Altace | - | - | - | - | 1/week | 42 |
| 9 (010) | Atorvastatin | - | - | Xanax, Zoloft, Wellbutrin, Imipramine | - | Spiriva | - | 0 | 45 |
| 10 (014) | Lipitor | Coreg, Lisinopril | Pradaxa | - | - | - | - | 0 | 80 |
| 11 (009) | - | Metoprolol, Enalapril | Lovenox | Lexapro, Buspar, Trazadone | Hydrocodone, Gabapentin | - | Type II (Metformin) | 0 | 82.5 |
| 12 (002) | - | - | - | Xanax | - | - | | 0 | 105 |
| 13 (001) | - | Hyzaar | - | Lexapro | - | - | Type II (Metformin) | 3/week | 20 |
| 14 (018) | - | - | - | - | - | - | | 2/day | 13 |
